# The Tissue Tropisms and Transstadial Transmission of a *Rickettsia* Endosymbiont in the Highland Midge, *Culicoides impunctatus* (Diptera: Ceratopogonidae)

**DOI:** 10.1101/2020.06.23.166496

**Authors:** Jack Pilgrim, Stefanos Siozios, Matthew Baylis, Gregory D. D. Hurst

## Abstract

*Rickettsia* are a group of intracellular bacteria which can manipulate host reproduction and alter sensitivity to natural enemy attack in a diverse range of arthropods. The maintenance of *Rickettsia* endosymbionts in insect populations can be achieved through both vertical and horizontal transmission routes. For example, the presence of the symbiont in the follicle cells and salivary glands of *Bemisia* whiteflies allows Belli group *Rickettsia* transmission via the germline and plants, respectively. However, the transmission routes of other *Rickettsia*, such as those in the Torix group of the genus, remain underexplored. Through fluorescence *in-situ* hybridisation (FISH) and transmission electron microscopy (TEM) screening, this study describes the pattern of Torix *Rickettsia* tissue tropisms in the highland midge, *Culicoides impunctatus* (Diptera: Ceratopogonidae). Of note is high intensity of infection of the ovarian suspensory ligament, suggestive of a novel germline targeting strategy. Additionally, localisation of the symbiont in tissues of several developmental stages suggests transstadial transmission is a major route of ensuring maintenance of *Rickettsia* within *C. impunctatus* populations. Aside from providing insights into transmission strategies, *Rickettsia* presence in the fat body of larvae indicates potential host fitness and vector capacity impacts to be investigated in the future.

**Importance Statement:** Microbial symbionts of disease vectors have garnered recent attention due to their ability to alter vectorial capacity. Their consideration as a means of arbovirus control depends on symbiont vertical transmission which leads to spread of the bacteria through a population. Previous work has identified a *Rickettsia* symbiont present in several vector species of biting midges (*Culicoides* spp.), however, symbiont transmission strategies and host effects remain underexplored. In this study, we describe the presence of *Rickettsia* in the ovarian suspensory ligament and the ovarian epithelial sheath of *Culicoides impunctatus*. Infection of these organs suggest the connective tissue surrounding developing eggs is important for ensuring vertical transmission of the symbiont in midges and possibly other insects. Additionally, our results indicate *Rickettsia* localisation in the fat body of *Culicoides impunctatus*. As viruses spread by midges often replicate in the fat body, this implies possible vector competence effects to be further investigated.

## Introduction

Heritable microbes of arthropods are important drivers of diverse host phenotypes. For example, both *Rickettsia* and *Wolbachia* are associated with reproductive parasitisms which favour the production of female offspring (e.g. male-killing and parthenogenesis) (1–3), whilst also being associated with resistance or tolerance against pathogens (4–6). Specifically, in disease vectors such as mosquitoes, both naturally occurring and artificially introduced symbionts can lead to a “virus-blocking” effect (7–10). These phenotypes, combined with maternal inheritance, drive the symbiont (and its effects on vectorial capacity) into a population and is currently being considered as a means of arbovirus control (11–13).

*Culicoides* biting midges (Diptera: Ceratopogonidae) are vectors which transmit economically important pathogens of livestock including bluetongue and Schmallenberg viruses (14). Currently, disease control primarily relies on vaccines which, given the rapid emergence and spread of these viruses, are often not available. Thus, alternate strategies, such as those based on symbionts, are of particular interest for midge-borne pathogens. So far, three endosymbionts have been observed in *Culicoides* spp.; *Wolbachia, Cardinium* and *Rickettsia* (15–18). Of these, symbioses with *Rickettsia* are the most common, and are present in several midge vector species (17). However, effects of the *Rickettsia* on the host are yet to be determined. The absence of sex-ratio distortion suggests the lack of a reproductive parasitism, and as some midge populations do not carry *Rickettsia* at fixation (excluding an obligate association), this indicates the drive of this endosymbiont might be related to a facultative benefit (e.g. pathogen protection).

This study focusses on broadening our understanding of the interactions between a Torix group *Rickettsia* and its host the highland midge, *Culicoides impunctatus*. The presence of *Rickettsia* in *C. impunctatus* oocytes has previously been identified and indicates transovariol transmission (17). However, the route of migration to egg chambers by the symbiont is not clear and the tropisms to other tissues remain unexplored. Vertical transmission of non-obligate symbionts is achieved through diverse modes of germline targeting (19). With certain *Wolbachia* strains of *Drosophila* spp., the symbiont localises in the germline stem cell niche continuously throughout development (20–22). For other symbionts, such as Belli and Adalia group *Rickettsia*, germline localisation follows infection of somatic tissues associated with the germline (e.g. follicle cells and bacteriocytes) (23–25). Intriguingly, *Rickettsia* also has the unusual ability to infect sperm head nuclei, allowing for paternal inheritance, which can combine with maternal transmission to drive a costly symbiont into the population (26). We therefore aimed to explore *Rickettsia* localisation in the midge germline and its associated tissues to glean insights into germline targeting mechanisms in this symbiosis.

Another objective of this project is to generate hypotheses of symbiont function through examining patterns of somatic tissue localisation. For instance, the close association of *Blattabacterium* with uric acid-containing cells (urocytes) in cockroaches is associated with nitrogen recycling into amino acids (27, 28), whilst the presence of *Wolbachia* in specific areas of the *Drosophila* brain is linked to mate choice (29). Furthermore, horizontal transmission pathways can be elucidated in a similar manner, with *Rickettsia* salivary gland infections of haematophagous and phloem-feeding arthropods reflecting transmission to vertebrates and plants respectively (24, 30, 31). In light of this, through fluorescence *in-situ* hybridisation (FISH) and transmission electron microscopy (TEM) screening, this study describes patterns of *Rickettsia* infection in both germline and somatic tissues in multiple developmental stages of *C. impunctatus*.

## Results

### Rickettsia infection during oogenesis

Early stage (stage 1) egg chambers contained clusters of *Rickettsia* predominantly in oocytes in which there is no presence of yolk deposition observed (Fig. 1A and 2A). The signal was strongest in the oocyte, with bacteria also being observed within nurse cells. Electron microscopy images suggested the *Rickettsia* are perinuclear rather than within the nurse cell nuclei themselves (Fig. 2B and C). The follicular epithelium was also infected with bacteria seen in transit between follicle cells and the oocyte (Fig. 1A). After 48 hours post-blood feeding, yolk deposition was seen as a clouding in the (stage 2) oocyte although individual yolk granules were not visible (Fig. 1B). Clusters of bacteria no longer occupied nurse cells but still predominantly filled the oocyte. In stage 4 eggs, yolk granules became visible and appeared to harbour sparse numbers of bacteria with a predominant localisation in follicle cells (Fig. 1D). However, in certain focal planes *Rickettsia* were observed in the oocyte cytoplasm but present only at the periphery of the oocyte (Fig. 1C).

**Figure 1.**
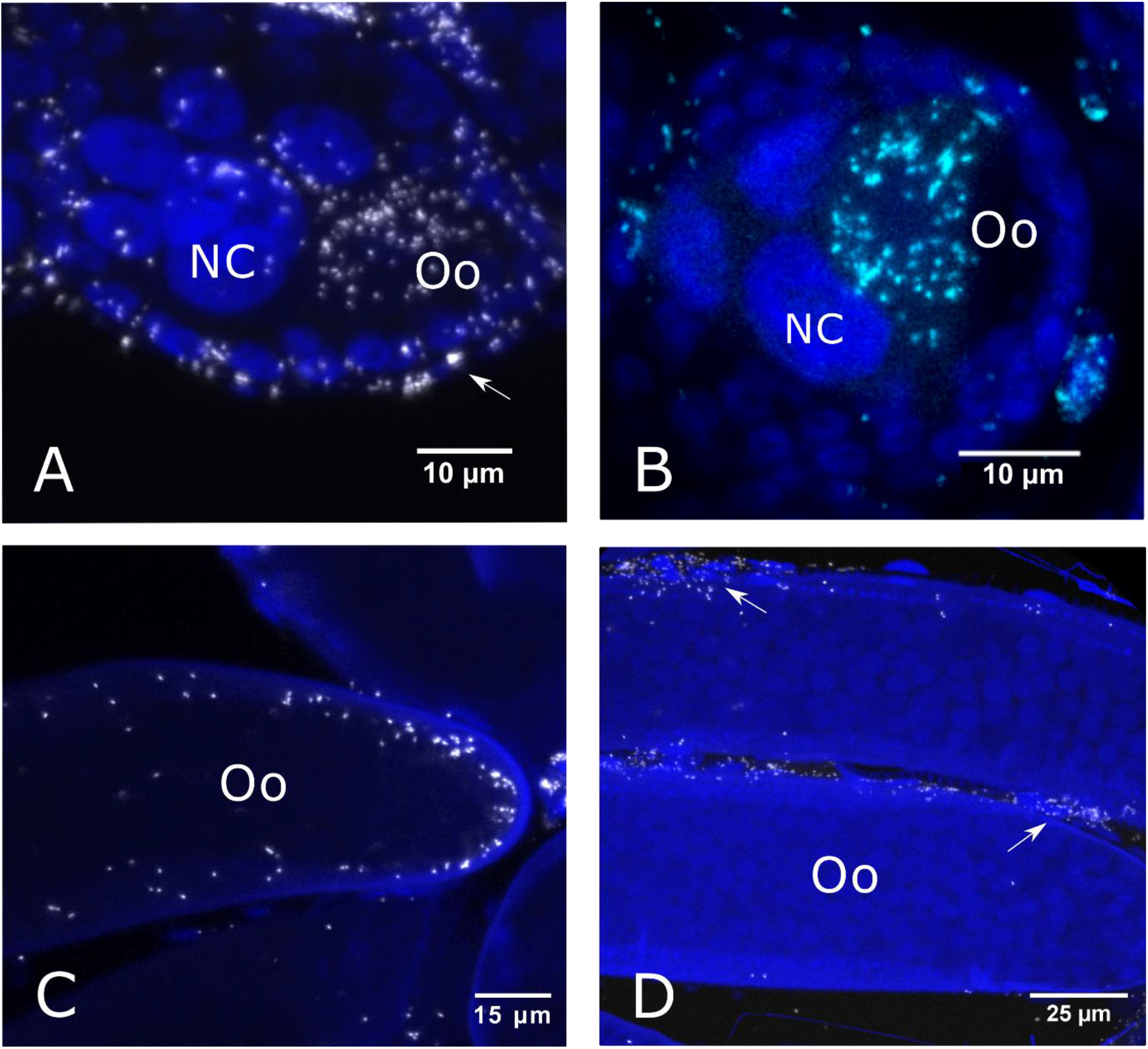
FISH images of *C. impunctatus* egg chambers at different developmental stages of oogenesis. *Rickettsia-specific* probe = white; DAPI-staining = blue. **A)** *Rickettsia* infection of stage 1 eggs (0 hours post blood-feeding) with predominant localisation in the Oocyte (Oo), Nurse cells (NC) and follicle cells (arrow). **B)** *Rickettsia* infection of stage 2 eggs (12 hours post blood-feeding) with the accumulation of cloudy yolk deposits in the oocyte (Oo). Infection is still primarily observed in the Oocyte (Oo) although infection of Nurse cells (NC) is now absent. **C)** A focal plane of stage 4 eggs (120 hours post blood-feeding) showing localisation at the periphery of the oocyte (Oo). **D)** A focal plane of stage 4 eggs (120 hours post blood-feeding) showing infection of follicle cells (arrows).

**Figure 2.**
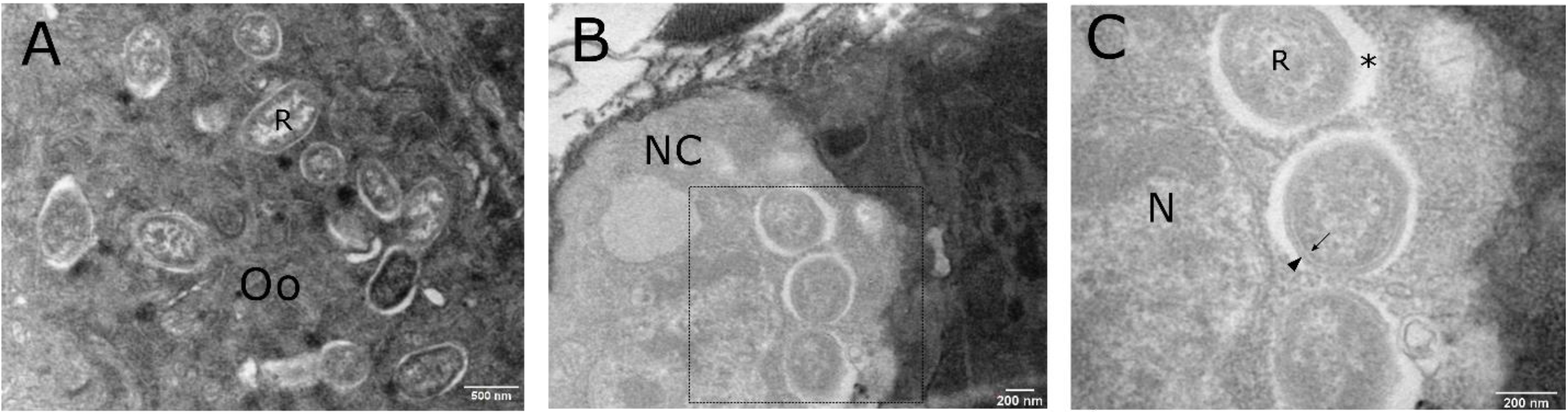
Transmission electron microscopy (TEM) images of *C. impunctatus* infected stage 1 egg chambers. **A)** TEM section of a *C. impunctatus* egg demonstrating clusters of *Rickettsia* in the oocyte cytoplasm (OO). **B)** TEM section of a *C. impunctatus* egg demonstrating *Rickettsia* presence in a nurse cell (NC). **C)** Magnified details of the box in C demonstrating perinuclear *Rickettsia* (R). The Rickettsiae have a distinctive cell wall (arrowhead) and cell membrane (arrow) separated by a periplasmic space. * = radiolucent halo/slime layer; N= nucleus.

### Rickettsia localisation to the ovarian suspensory ligament

Initial examination of adult specimens at low magnification gave a consistent pattern of strong localised signal at the anterior/posterior midgut junction (Fig. 3A). On examination at a higher magnification, no signal was observed in any of the midgut epithelial cells (Fig. 3C). Further scrutiny of individuals led to the discovery that the infected structure was the ovarian suspensory ligament. This structure, otherwise known as the “median ligament”, was seen to pair off and loop down from the midgut attachment site before attaching at the apex of the ovary (Fig. 3A to C). It was possible to follow the signal down the suspensory ligament where the structure became continuous with the terminal filaments of ovarioles and the ovarian epithelial sheath (Fig. 3B and D), the structure encasing ovarioles. Strength of infection was consistent in the ovarian epithelial sheath over the long axis of the ovary with neighbouring immature egg chambers seen to be heavily infected. Bacteria could be seen migrating from this densely populated structure into neighbouring follicle cells and further into oocytes themselves (Fig. 3D). Within the ovary, the germarium appeared to be no more strongly infected than the rest of the ovarian tissue. In one individual, the attachment of the suspensory ligament to a sparsely cellular but strongly signalled structure is thought to be part of a lobe of the fat body (Fig. 3A) although the ethanol based (Carnoy’s) fixative diminishes lipids leading to ambiguity of identification when observed under transmission light.

**Figure 3.**
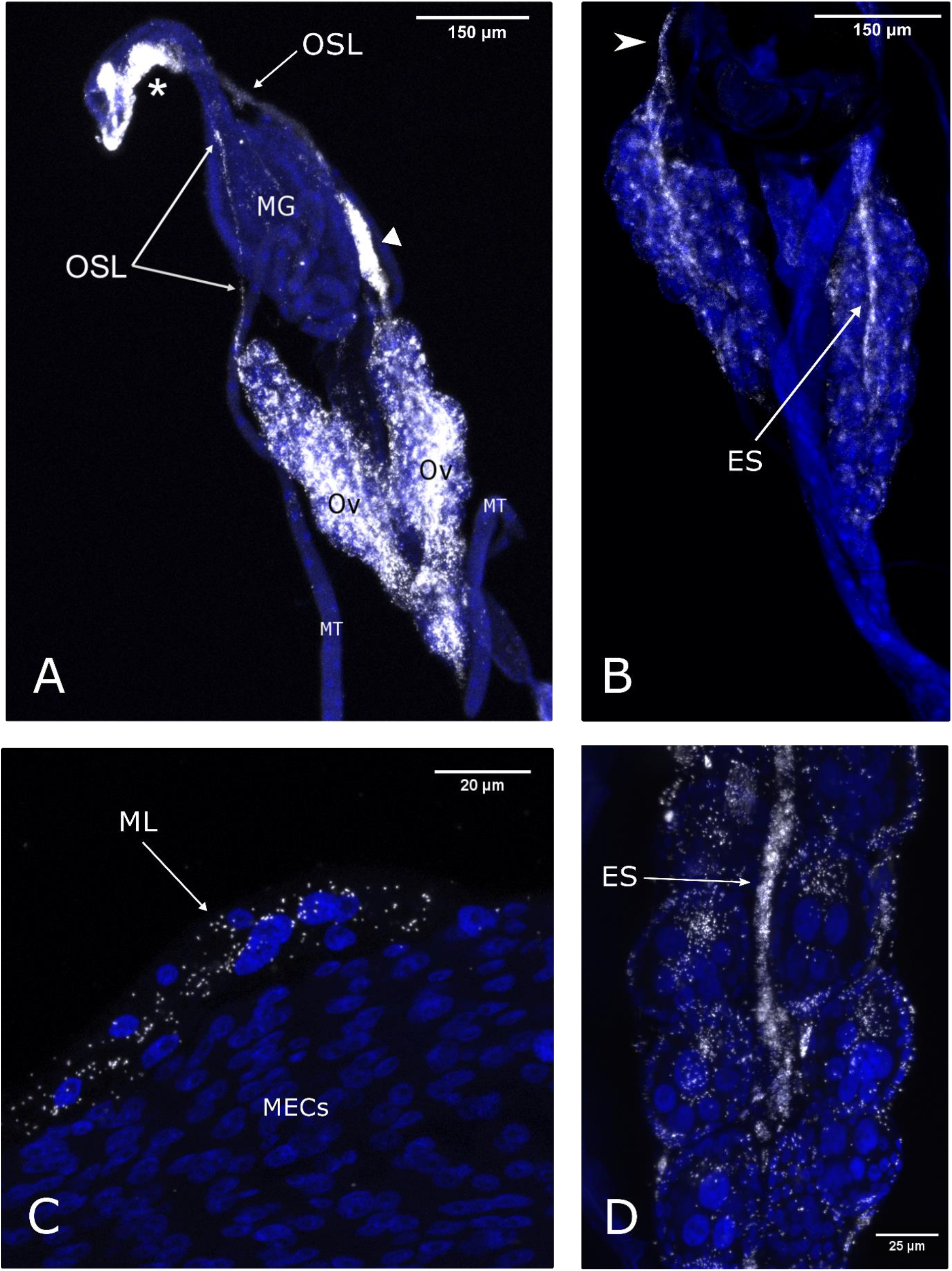
*Rickettsia* localisation in *C. impunctatus* adult connective tissues associated with both the midgut and ovaries via FISH imaging. *Rickettsia-specific* probe = white; DAPI-staining = blue. **A)** Strong *Rickettsia* signals identified at the anterior-posterior midgut junction (*) as well as the paired ovaries (Ov). These two areas are connected via the ovarian suspensory ligament (OSL) which runs from the midgut junction to the apex of the ovary. White triangle=putative fat body lobe; MG=midgut; MT=Malphigian tubules. **B)** A focal plane of the paired ovaries demonstrating the continuation of the suspensory ligament attachment site at the ovary apex (arrowhead) into the ovary. **C)** *Rickettsia* localisation at the median ligament (ML); the fusion of the ovarian suspensory ligaments at the attachment site at the anterior-midgut junction. Lack of infection is observed in midgut epithelial cells (MECs). **D)** The continuation of the ovarian suspensory ligament with the ovarian epithelial sheath (ES) allows for the delivery of *Rickettsia* into neighbouring egg chambers.

### Rickettsia infection in other adult tissues

In the single male specimen available for analysis, infection of the testes was observed (Supplementary Fig. 1B). *Rickettsia* was further detected in crushed spermathecae from fertilised females (Fig. 4A). Subsequently, TEM sections of spermathecae were assessed to clarify the nature of this signal. Overall, infected spermathecae showed no evidence of bacteria in sperm heads or tails of spermatids, nor in the acellular matrix (Fig. 4B). However, *Rickettsia* was identified in the maternally-derived spermathecal epithelium (Fig. 4C and D). Unfortunately, due to the difficulties of laboratory maintenance of *Culicoides*, a crossing-system to definitively rule out paternal transmission was not possible. Finally, a single infection was observed in the crop of the foregut (Fig. S1A) with no signal observed in Malpighian tubules, heads or salivary glands for all adult samples.

**Figure 4.**
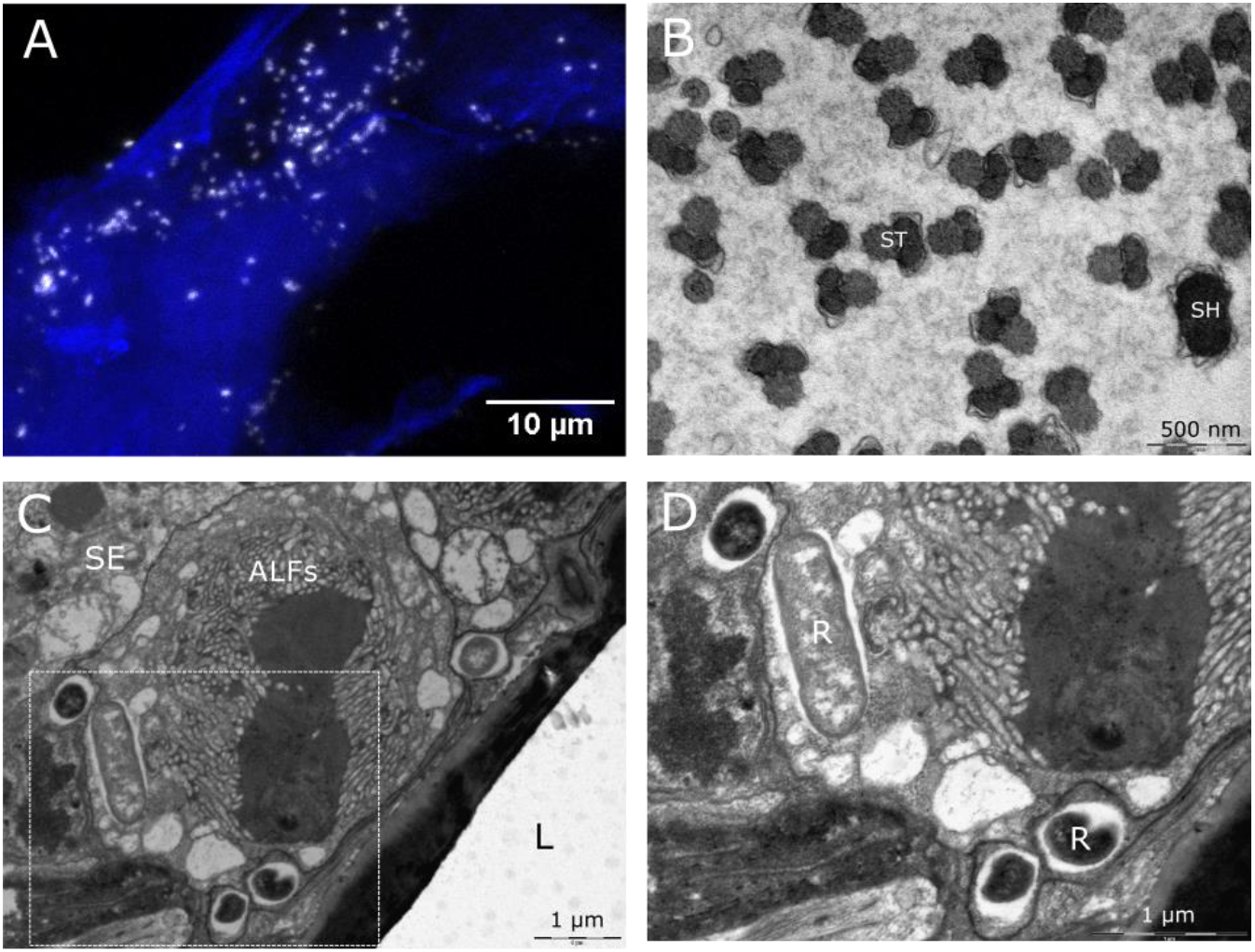
FISH and TEM analysis of *C. impunctatus* spermathecae. **A)** FISH image of *Rickettsia* presence in a crushed spermatheca. *Rickettsia-specific* probe = white; DAPI-staining = blue. **B)** TEM section of *Rickettsia* uninfected sperm heads (SH) and sperm tails (ST). **C)** TEM section of junction between the spermathecal lumen (L) and spermathecal epithelium (SE). ALFs=Actin-like filaments. **D)** Higher magnification details of the box in C, demonstrating longitudinal and cross-sectioned *Rickettsia* (R) residing in the spermathecal epithelium.

### Subcellular location and associations

Transmission electron microscopy of spermathecae and immature eggs revealed coccobacilli *Rickettsia* free in the cytoplasm of oocytes and spermathecal epithelial cells (Fig. 2A and 4C to D). Sections of bacteria ranged up to 1.35 μm in length and were seen together either in clumps, likely a result of recent division, or diffusely in tissues. The ultrastructure of *Rickettsia* demonstrated distinctive characteristics typical of the genus; a slime layer/radiolucent halo, and an outer trilaminar cell wall followed internally by a periplasmic space and membrane (Fig. 2C). Infection was not observed in host cell nuclei.

### Larval tissue localisation

Out of ten L3 larvae of unknown infection status, four showed a positive signal in the terminal abdominal (anal) segment fat body (Fig. 5B and 6A) with three of these also demonstrating infection in the heads (Fig. 6B). One individual of the four positives showed sporadic multifocal infections across the rest of the length of the body. Focal “burst” patterns of signal in the terminal abdominal segment suggest infections of globe-like structures such as cells or lipid droplets of the fat body. Infections in the head could not be localised to the exact tissue infected, but *Rickettsia* were closely associated with the head body wall (within 0-3 μm proximity of the autofluorescent cuticle of each focal plane). Concurrent presence in the large fat bodied anal segment of the larvae, alongside the fat body’s frequent attachment to the body wall suggests the pericerebral fat body is the most likely tissue to be infected, although further work is needed to confirm this.

**Figure 5.**
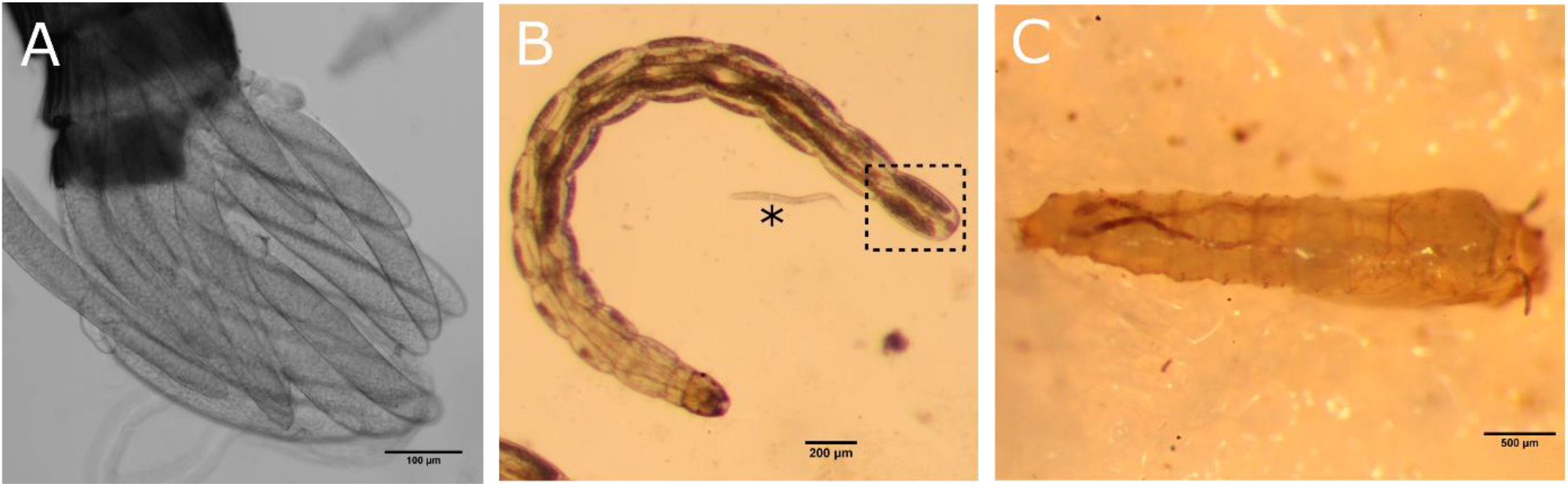
Transmitted light microscope images of different life stages of *C. impunctatus*. **A)** Stage 4 eggs. **B)** L3 larva with lateral fat bodies terminating in the terminal abdominal segment (Box); * = *Panagrellus nephenticola* nematode. **C)** Pupa.

**Figure 6.**
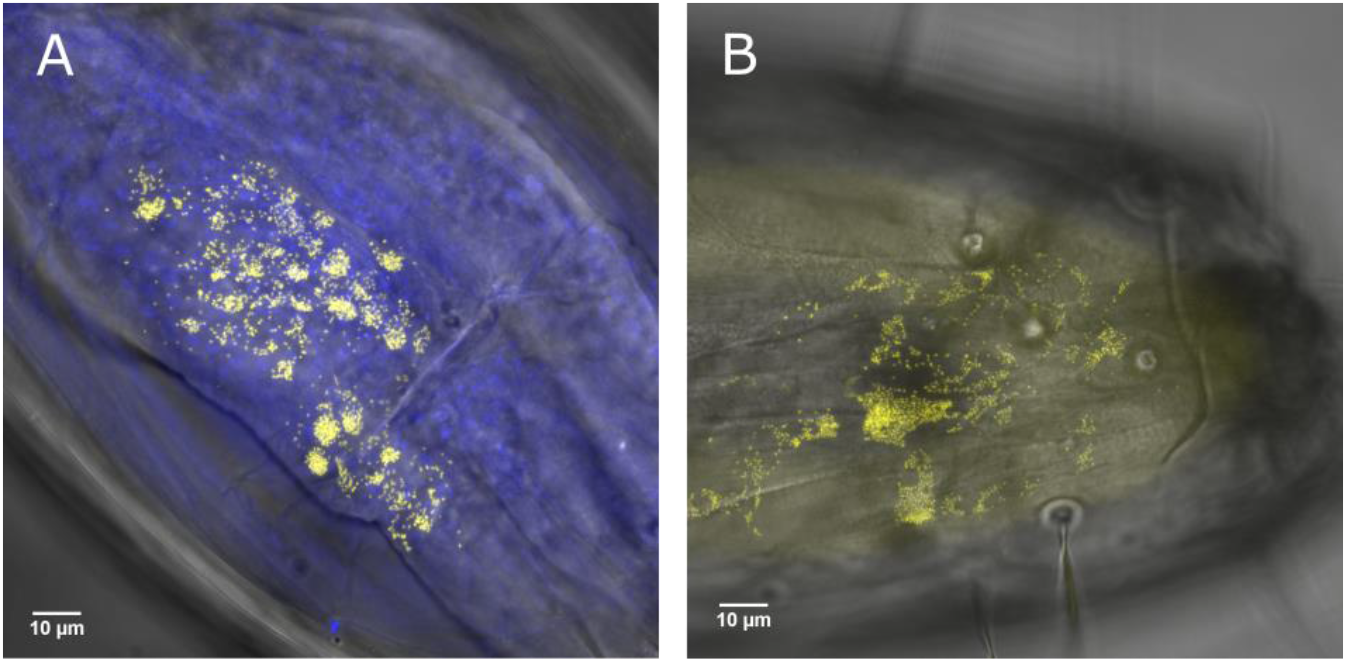
FISH imaging analysis of an L3 *C. impunctatus* larva. *Rickettsia-specific* probe = yellow; DAPI-staining = blue. **A)** Fat body *Rickettsia* infection of the terminal abdominal segment. **B)** *Rickettsia* larval head infection.

## Discussion

*Rickettsia* bacteria are important components of arthropod biology contributing to host protection against pathogens (5, 6), whilst also being causative agents of disease in their own right (32). The transmission routes of *Rickettsia* endosymbionts in biting midges are underexplored but can be informed by symbiont tissue localisation. Although the maternal transmission of obligate symbionts are generally dependent on specialised cells (bacteriocytes), facultative (secondary) symbionts utilise various means to target the germline (19). These can come in the form of co-option of yolk granules to gain entry into egg chambers via endocytosis (e.g. *Spiroplasma* in *Drosophila melanogaster*) (33), or through the continuous association with the germline during morphogenesis (e.g. *Wolbachia* in *Drosophila* spp.) (Veneti *et al*., 2004; Serbus and Sullivan, 2007). These modes of germline targeting can be dismissed in the case of *C. impunctatus*-*Rickettsia* symbioses. Oocyte symbiont infection precedes yolk deposition, indicating no route through cooption of the vitellogenin transport systems (Fig. 1A), and larval stages show no localisation to the mid abdominal area in which the germline progenitors are present (infection is in the head and terminal segments) (Fig. 6). Intriguingly, our description of the *Rickettsia* infected ovarian suspensory ligament in *C. impunctatus*, indicates a potential novel means of endosymbiont germline targeting (Fig. 3).

The continuation of the suspensory ligament with the ovarian epithelial sheath (Fig. 3B and D) has previously been described in insects (34), with our results suggesting these tissues act as an intermediary for ensuring *Rickettsia* infection of egg chambers via follicle cells. The passage of *Rickettsia* through the follicle cells of ovarioles occurs in the ladybird *Adalia bipunctata* and the whitefly *Bemisia tabaci* (23, 24). In these instances, it is possible that *Rickettsia* migrates through follicle cells after ovarian contact with infected hemocytes/hemolymph (24, 35). Other studies have suggested bacteriocytes could be a means of transovarial transmission for *Rickettsia* (25, 36, 37), although further investigation suggested this was not a primary route of ovary-targeting in whiteflies (24). In light of this, it is conceivable that infection of connective tissue directly linking the germline allows for the vertical transmission of *Rickettsia* in midges and other insects.

Our findings of *Rickettsia* infection in follicle cells but with apparent limited infection of mature oocytes (Fig. 1C and D), corroborates previous observations in the whitefly *Bemisia tabaci* (24). In this case, heavy *Rickettsia* infection in immature oocytes, but not mature stages, was attributed to younger eggs being more permeable than their mature counterparts. Alternatively, it may be the case that this perceived density variation is the result of a dilution effect as the oocyte gets bigger. The *Rickettsia* infection and close contact of the ovarian epithelial sheath to the follicle cells offers a mechanism for ensuring persistent follicular infection (Fig. 3D). Furthermore, as only a few bacteria cells are required to ensure subsequent infection of life stages (26), remnant *Rickettsia* in the oocyte periphery of mature eggs (Fig. 1C) appears to be sufficient for transstadial transmission (Fig. 6).

Although intranuclear infections are rare for bacteria, various *Rickettsia* strains have been observed to reside within nuclei (26, 37–39) with no indication of a detrimental impact on arthropod reproduction or development. Of particular interest is a Torix group *Rickettsia* which undergoes paternal transmission via sperm head nuclei in the leafhopper *Nephotettix cincticeps* (26). The additive effects of maternal and paternal transmission not only solidify vertical transmission but can also drive symbiont evolution through lineage mixing in the absence of any benefit of infection. In the current study, initial localisation of *Rickettsia* to testes and spermathecae (Fig. 4A and 1B) suggested intrasperm transmission may also be occurring in *C. impunctatus*. However, TEM sections of spermathecae provided evidence only for *Rickettsia* infection of maternally-derived epithelia with no bacteria observed in sperm head nuclei. Similarly, nurse cells of ovaries demonstrated the presence of perinuclear *Rickettsia* only (Fig. 2C). The absence of an intranuclear tropism in eggs as well as sperm suggests the capacity to infect nuclei may be evolutionary labile.

While vertical transmission is the primary transmission route for endosymbionts, horizontal transfer frequently occurs in *Rickettsia* and has allowed for the infection of a diverse range of organisms including protists, arthropods, plants and vertebrates (40, 41). The lack of a *Rickettsia* infection signal in salivary glands of *C. impunctatus*, and the parity of prevalence in male and female midges (17) indicates horizontal transmission to vertebrate hosts via haematophagy is unlikely.

Another finding of interest is the localisation of *Rickettsia* to the fat bodies of larvae (Fig. 6A). Previous examples of endosymbiont fat body infections of *Culicoides* include the observation of “Rickettsia-like” organisms by Hertig and Wolbach (42) and “twinkling” symbionts under polarised light by Lawson (43). The larval fat body largely comprises two bands extending down the lateral abdomen terminating in large lobes of the terminal segment (43, Fig. 5B). *Rickettsia* appears to be one of few known endosymbionts to reside in the fat body with others including *Blattabacterium* of cockroaches (44) and *Wolbachia* in a variety of insects (45–47). The presence of *Wolbachia* in the fat body, an important endocrine tissue, has been associated with effects on host glucose and glycogen metabolism via altered enzyme activity and insulin signalling (48, 49). Not only is the fat body metabolically active, allowing for the bacterial sequestering of metabolite precursors, but fat cells are refractive to degradation during larval and pupal development (50, 51). Thus, this tissue offers a suitable niche in arthropods for stable division and proliferation of endosymbionts.

Other *Rickettsia* infected regions of note are larval heads (Fig. 6B). Although it is not clear exactly which tissues are affected by the *Rickettsia*, the pericerebral fat body and cerebral ganglia (brain) are two candidates for future consideration. Endosymbiont brain infections have been proposed to lead to behavioural modifications (29, 52). Additionally, the fat body surrounding the brain has been demonstrated to play a different physiological role to that in the abdomen. Namely, unique insulin signalling pathways have been observed in the head fat bodies of *Drosophila melanogaster* leading to increased longevity as a result of inhibited senescence (53).

The presence of *Rickettsia* in the fat body of *Culicoides*, a vector of several veterinary viruses, raises questions on their effects on vector competence. Bluetongue virus (BTV) and epizootic haemorrhagic disease (EHDV) of ruminants replicate in the fat body of midges (54, 55) before travelling to the salivary glands, suggesting interactions between *Rickettsia* and the virus could be occurring. For example, competition for lipids between bacteria and virus have been suggested to influence viral titres (56–59). Additionally, antimicrobial peptides are synthesised in the fat body (60) and are active against arboviruses (61), again suggesting *Rickettsia* effects on vectorial capacity warrant further investigation.

*Culicoides impunctatus* are a biting nuisance in the Scottish Highlands, with “midge attacks” accountable for significant economic impact through losses in tourist and forestry industries (62, 63). Their ability to reproduce once in the absence of a blood meal (autogeny) means huge numbers can develop even where vertebrate hosts are not available (64–66). Thus, it is interesting to speculate that *Rickettsia* may play a possible role in autogeny. Indeed, *Rickettsia felis* is necessary for egg development in the booklouse *Liposcelis bostrychophila* (67). As autogeny is responsible for the pest burden of *C. impunctatus*, this could offer a target for population suppression in the future. Unfortunately, the difficulties in maintaining *C. impunctatus* colonies is a hinderance in investigating such host effects. The rearing methods of this study allowed for development to pupae (Fig. 5C) but overall survival was poor with none completing a full life cycle (Fig. S2). Thus, further optimisation of *C. impunctatus* rearing is needed to investigate this symbiotic system further.

In conclusion, this study has identified several somatic and germline infections of a Torix *Rickettsia* endosymbiont in several biting midge life stages. Infection of the ovarian suspensory ligament, a continuation of the ovarian epithelial sheath, has identified a potential novel means of endosymbiont germline targeting. Additionally, a somatic tissue infection to study further is the fat body, which could have implications for host effects, as well as arbovirus transmission dynamics.

## Materials and methods

### Culicoides impunctatus collections

Collections of *Culicoides impunctatus* for imaging of adult tissues were carried out at multiple sites in the United Kingdom; Dumfries, Kielder forest, Loch Lomond and Fort William between June 2017 and September 2018. For midges not used for ovipositing, individual *Culicoides* were allowed to rest on an author’s arm (JP + SS) before aspiration into 1.5 ml Eppendorf tubes with 10% sucrose-soaked cotton wool placed at the bottom and damp sphagnum moss (B&Q, UK) filled in the lid to maintain a high humidity. Tubes were then placed horizontally on sticky tape in plastic boxes before incubation at 23 °C. For ovipositing *Culicoides*, female midges were collected by allowing feeding on an author (JP + SS) and field assistants’ forearms from the Loch Lomond site (May 2018). When midges were observed to be replete (approximately after 5 minutes), or the insect released mouth parts from the skin, they were aspirated and stored in a plastic container at ambient temperature (13-17°C) until the end of the collection session.

Oviposition containers were assembled based on previous studies undertaken by Carpenter (68). Briefly, approximately 50 *Culicoides* were transferred to containers consisting of cylindrical pill boxes (Watkins and Doncaster, UK; 64 mm diameter x 60mm depth), with cotton wool soaked in 10% w/v sucrose solution (replaced every two days) placed on top of a fine net meshing which covered the tops of the pillboxes. For ovipositing areas, 50 ml Falcon tube lids were filled with damp sphagnum moss by soaking in 1% nipagin dissolved in distilled water and squeezing until drops could be counted, before being placed on top of damp filter paper. This was secured by cutting a cylindrical hole in the bottom of the pill box. These were then transferred to 19 x 12 x 8 cm plastic boxes containing a 50 ml beaker of saturated sodium sulphate (Na_2_SO_4_) which has previously been described to maintain humidity at >90% when between 20-25°C (69). These were then transported to a laboratory in Liverpool before the plastic boxes were placed in an incubator where they were maintained at 23°C with a photoperiod of 12 Light: 12 Dark hours.

### Larval rearing

Eggs oviposited onto the sphagnum moss substrate were picked individually with a fine paintbrush or the damp edge of a sharpened tungsten needle and placed onto 0.5% agar dishes (100-180 eggs per dish; n=3 dishes) and spaced evenly apart. *Culicoides impunctatus* identification was confirmed by the distinctive brown heads of larvae (70) as well as identification of ovipositing adults in pill boxes. Larvae were fed daily on Banana worms (*Panagrellus nepenthicola*) which were cultured using the manufacturer’s instructions (Ron’s worms, UK) before a fine paintbrush was used to place the nematodes in 2-3 ml of deionised water and spread evenly across each agar dish. Immatures were stored under the same temperature and photoperiods as adults previously mentioned. Allocation of larval instars were designated by head capsule length measurements as described by Kettle and Lawson (70).

### Dissections of adults

Post blood-feeding in the wild, *Culicoides* were processed after carefully timed transportation to a laboratory in Liverpool. Individuals were sacrificed at various time points post blood feeding (PBF); 0 hours (non-blood fed), 12 hours PBF, and 120 hours PBF. First, *Culicoides* were chilled in the freezer for immobilisation before being placed in a drop of Phosphate-buffered saline on a petri dish. Midges were then killed by piercing the thorax with a sharp tungsten needle before being confirmed as *C. impunctatus*. Ovaries and other tissues were then exposed through dissection under a stereoscopic microscope. Time points for sacrifice were informed by Carpenter (68) which identified developmental stages of forming eggs as a function of time after blood-feeding. Stages were confirmed by using a system developed by Linley (71) which was subsequently modified by Campbell and Kettle (72):

stage 1) No observation of yolk within the oocyte;
stage 2) Yolk can be identified within the oocyte;
stage 3) Yolk proteins occupy up to three-quarters of the oocyte;
stage 4) The oocyte is elongated and no longer oval, resembling the mature egg;
stage 5) Egg fully mature, with chorion visible.

### Tissue preparation and fluorescence in-situ hybridisation (FISH)

Tissues examined included eggs of different developmental stage, Malpighian tubules, midgut, foregut, hindgut, fat body, testes and salivary glands. Additionally, crushed spermathecae were prepared for visualisation of the spermatophore and spermatids contained within. To this end, spermathecae were suspended in Phosphate-buffered saline and allowed to dry before pressing a coverslip over the slide to break open the tissue.

For fluorescence *in-situ* hybridisation (FISH) imaging, the above tissues were fixed directly on poly-L-lysine covered slides for one hour in Carnoy’s solution (chloroform:ethanol:glacial acetic acid, 6:3:1) and tissues cleared by treating with 6% H_2_O_2_ in ethanol for 2 hours. Two pre-hybridisation washes were undertaken using wash buffer (20 mM Tris-HCl, pH 8.0, 50mM NaCl, 0.01% sodium dodecyl sulphate, 5 mM EDTA). Hybridisation was performed overnight in hybridisation buffer (20 mM Tris-HCl, pH 8.0, 90mM NaCl, 0.01% sodium dodecyl sulphate, 30% formamide) containing 10 pmol/ml of the *Rickettsia* specific probe [5’-CCATCATCCCCTACTACA- (ATTO 633)-3’] adapted from Perotti *et al*. (67) which were checked for specificity against the Torix 16S gene of *Culicoides newsteadi* (MWZE00000000). After hybridisation, the samples were thoroughly washed twice in wash buffer and slide mounted in Vectashield with DAPI (Vector Laboratories) and viewed under a Zeiss LSM 880 BioAFM confocal microscope. *Rickettsia*-free midges (*Culicoides nubeculosus*; Pirbright Institute, UK) were used as negative controls. For each tissue, at least 5 specimens were viewed under the microscope to confirm reproducibility (except a single adult male available for analysis). Optical sections (0.7μm thick) were prepared from each specimen to create a Z-stack image to be processed in ImageJ. All FISH imaging equipment and technical assistance was provided by the Liverpool Centre for Cell Imaging (University of Liverpool, UK).

### Transmission electron microscopy

As host-seeking *Culicoides* mate prior to blood feeding it was possible to examine spermatids from female spermathecae. Ovaries and spermathecae were prepared for transmission electron microscopy (TEM) as follows: Tissues were dissected into 2% (w/v) paraformaldehyde + 2.5% (w/v) glutaraldehyde in 0.1M phosphate buffer (pH 7.4). Fixative was then changed for 2.5% (w/v) glutaraldehyde in 0.1M phosphate buffer (pH 7.4). Heavy metal staining consisted of 2% (w/v) OsO4 in ddH_2_O, followed by 1% (w/v) Tannic acid in ddH_2_O and then 1% (w/v) aqueous uranyl acetate. To prevent precipitation artefacts the tissue was washed copiously with ddH_2_O between each staining step. Fixation and staining steps were performed in a Pelco Biowave^®^Pro (Ted Pella Inc.Redding California, USA) at 100W 20Hg, for 3 mins and 1 min respectively. Dehydration was in a graded ethanol series before filtration and embedding in medium premix resin (TAAB, Reading, UK). For TEM, 70-74 nm serial sections were cut using a UC6 ultra microtome (Leica Microsystems, Wetzlar, Germany) and collected on Formvar (0.25% (w/v) in chloroform, TAAB, Reading, UK) coated Gilder 200 mesh copper grids (GG017/C, TAAB, Reading, UK). Images were acquired on a 120 kV Tecnai G2 Spirit BioTWIN (FEI, Hillsboro, Oregon, USA) using a MegaView III camera and analySIS software (Olympus, Germany).

## Declarations

## Acknowledgments

We would like to thank Dr. Ewa Chrostek for kindly providing comments on the manuscript. We also thank Charlie Winstanley, Matthew Palmer and Lukasz Lukomski for their support with the collection of midge samples. This work was supported by a BBSRC Doctoral Training Partnership studentship awarded to JP. We acknowledge the Liverpool Centre for Cell Imaging (CCI) for provision of imaging equipment and technical assistance (BB/M012441/1). The processing of samples and electron microscopy imaging was undertaken with assistance from Alison Beckett and the Electron Microscopy Unit (Institute of Translational Medicine; University of Liverpool). *Culicoides nubeculosus* were provided via a Core Capability BBSRC Grant awarded to Simon Carpenter (The Pirbright Institute) and produced by Eric Denison (BBS/E/I/00001701).

## Author contributions

Field and Laboratory work was undertaken by JP and SS. Analysis and interpretation of the data were undertaken by JP, SS and GH, as well as drafting of the manuscript. GH and MB assisted in the conception and design of the study, in addition to critical revision of the manuscript.

