## Supplemental Figures 1 and 2 for "The Tissue Tropisms and Transstadial Transmission of a *Rickettsia* Endosymbiont in the Highland Midge, *Culicoides impunctatus* (Diptera: Ceratopogonidae)"

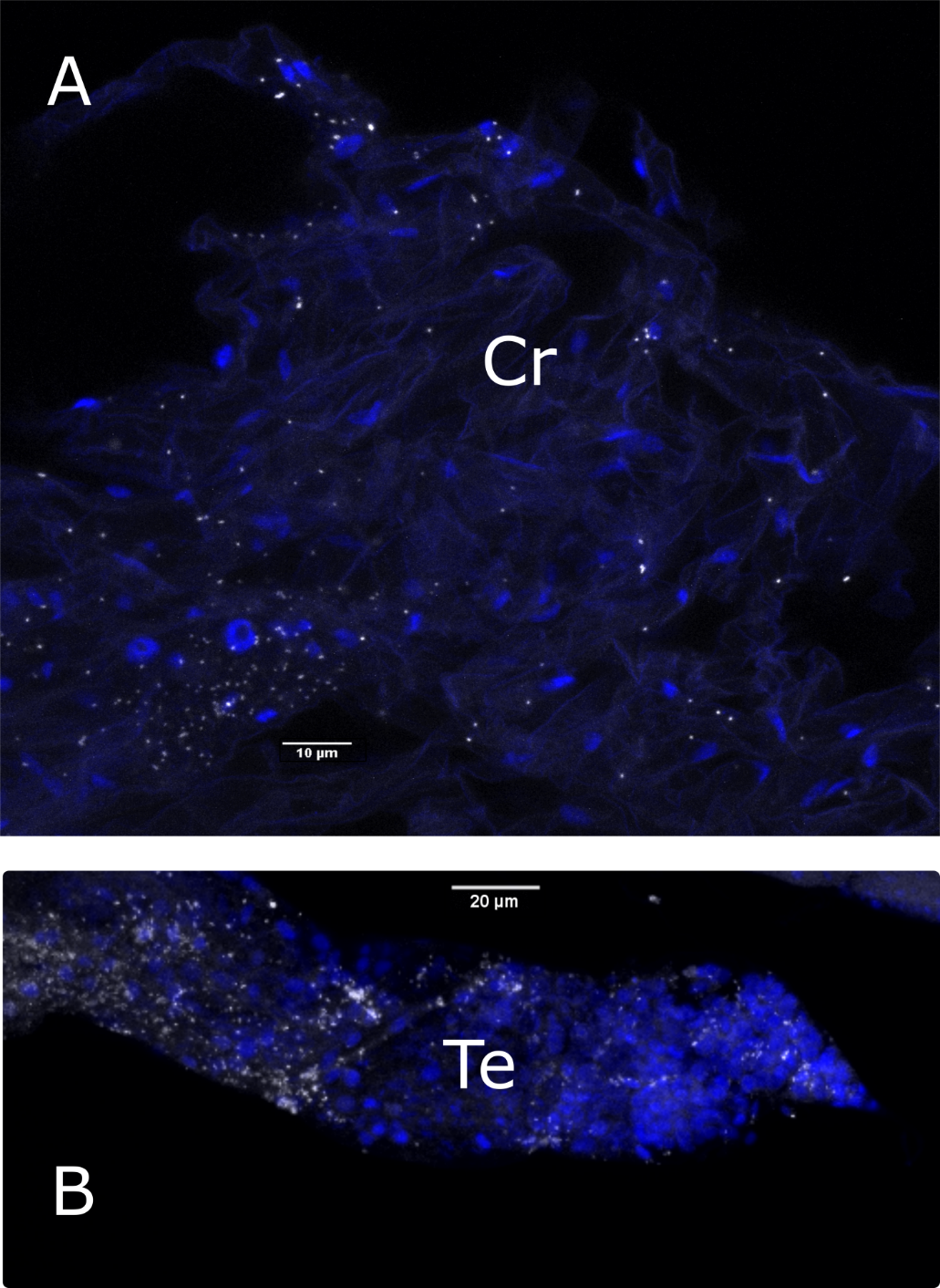

**Supplementary Figure 1.** *Rickettsia* infections of **A)** crop (Cr) and **B)** testes (Te). *Rickettsia*-specific probe = white; DAPI-staining = blue.

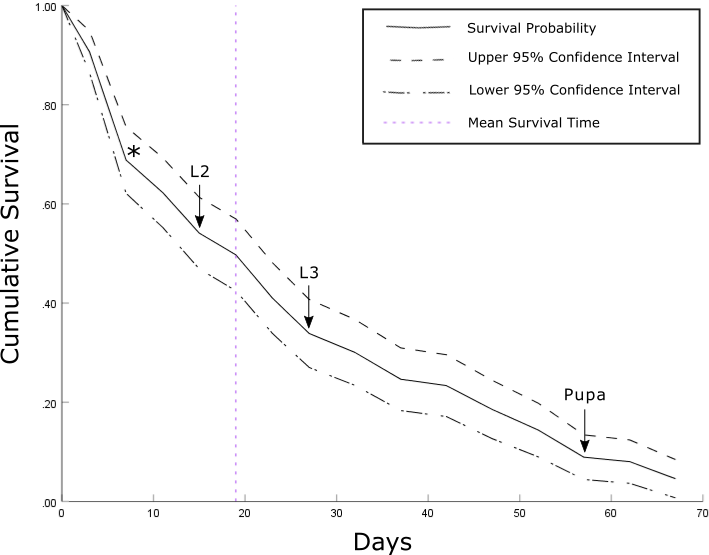

**Supplementary Figure 2.** Kaplan-Meier survival curve monitoring cumulative survival of *C. impunctatus* larvae (n=183) over time. Arrows demonstrate the first appearance of different instars. The asterisk is the point at which cannibalism ceased to be observed in L1s. Development into pupae (n=5) took a minimum of 56 days and the mean survival time of larvae was 19 days. **NB**-the burrowing behaviour of mature larvae made for difficult retrieval and head measurements, meaning a formal identification of the first appearance of L4 instars was not achieved.
